# cNPDB: A comprehensive empirical crustacean neuropeptide database

**DOI:** 10.1101/2025.07.29.667494

**Authors:** Vu Ngoc Huong Tran, Thao U. Duong, Lauren Fields, Kosta Tourlouskis, Margot Beaver, Lingjun Li

## Abstract

Neuropeptides, key signaling molecules essential for dynamic regulation of biological processes, have been studied via crustacean model systems for more than 40 years. We present cNPDB, the first centralized resource dedicated to this dynamic area of research, promisingly accelerating crustacean neuropeptidome research and offering a blueprint for cross-species translational studies of neuropeptide signaling. cNPDB is comprised of 1364 published neuropeptides from 29 crustacean species, each annotated with corresponding taxonomic information, computed physicochemical properties, spatial localization, and predicted three-dimensional structure. The intuitive web-interface supports keyword and property-based filtering, sequence alignment, property calculation, and links to available mass spectrometry imaging data, streamlining discovery.

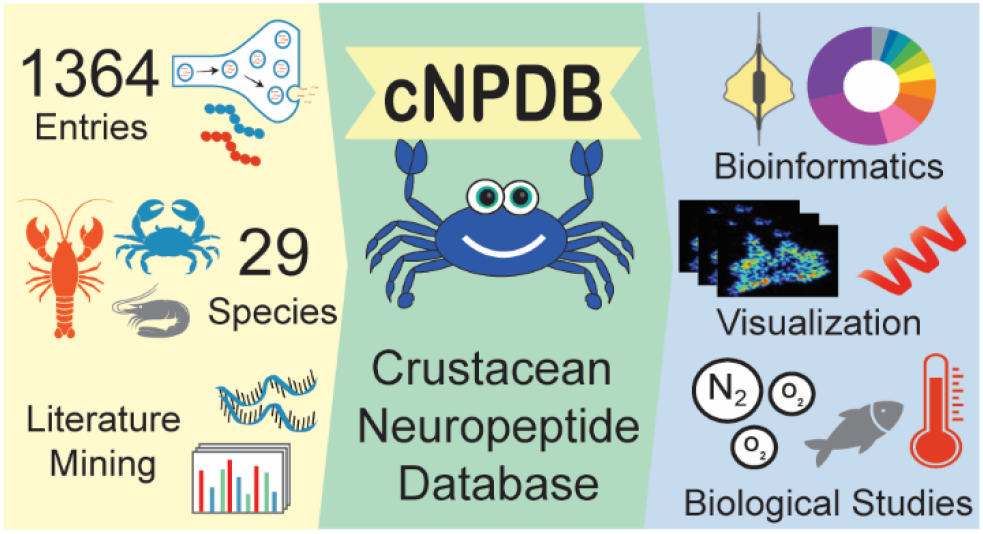

## Introduction

Neuropeptides are the most structurally diverse and functionally versatile signaling molecules, which play pivotal roles in many physiological processes, including appetite control, stress response, reward processing, and pain modulation.(1-5) Differing from classical neurotransmitters, neuropeptides can act over longer distances and durations, with potent, selective effects in the nervous system at nanomolar concentrations. Because of their precise and long-lasting actions, neuropeptides are promising candidates for developing targeted therapies and diagnostic tools for neurological and psychiatric disorders.(6-12) Despite their potential, neuropeptides remain challenging to characterize due to their high sequence variability, extensive post-translational modifications (PTMs), low endogenous concentrations, and rapid degradation. In mammalian nervous systems, with layered neural co-modulation, these complexities are further amplified, deepening the burdens associated with functional neuropeptidomics.(13) Invertebrate models, including crustacea, have simpler neural architectures and well-mapped peptidergic systems, which provide powerful platforms for uncovering conserved neurobiological mechanisms relevant to mammals.(14,15) Many neuropeptide families, defined by conservation of function and sequence, have established homologs between invertebrates and mammals.(16)

To date, four major online databases have been developed to support neuropeptide research, each with their own strengths and limitations, though none are exclusive or encompassing of the crustacea (**Table 1**). Focused on neuropeptides encoded by mammalian and vertebrate genes, www.neuropeptides.nl was the first online resource available, published in 2010.(17) NeuroPedia, released in 2011, centered on neuropeptides within phylum *Chordata*, excluding arthropods.(18) We should note that, to our knowledge, NeuroPedia is no longer accessible. Also exclusive of crustacea, DINeR is limited to species in the class *Insecta* only.(22) While NeuroPep covers both vertebrate and invertebrate species to facilitate drug design and new neuropeptide identification, its curation did not focus on the subphylum *Crustacea*, leaving many known crustacean neuropeptides unreported.(19,20) *Crustacea* collectively have advanced our understanding of environmental adaptation,(23-26) development and reproduction,(27-29) feeding,(30,31) immunomodulation,(32-35) and hypoxia,(36,37) impacting the diverse fields of endocrinology, genomics, neuroscience, physiology, and beyond. Additionally, fundamental studies from crustacea have elucidated the dynamic impact of PTMs,(38-41) peptide isoforms,(42,43) D-amino acid containing peptides,(44-46) and more at the molecular level, providing insights into rhythmic behaviors(47) and their underlying oscillatory networks,(48,49) neural remodeling,(50) and gene interaction.(51,52) Given these broad contributions, the absence of centralized crustacean database for the community is a hindrance to research progress. Herein, we describe our new database resource, Crustacean Neuropeptide Database (cNPDB), freely available online at https://lilabs.org/cnpdb, for researchers to reference (**Figure 1**).

**Table 1:** Database comparisons between neuropeptides.nl, NeuroPedia, NeuroPep 2.0, DINeR, and cNPDB in 4 categories: year of introduction, coverage, key data/ features, and data sources.

| Database Name | Year | Coverage | Key Data/Features | Sources |
| --- | --- | --- | --- | --- |
| **Neuropeptides.nl**<br>(17) | 2010 | Mammalian,<br>Vertebrate | 279 neuropeptides, 115 precursors, 17 gene families, over 95 gene names, mouse brain expression. Provides a historical overview, neuropeptide definition, and hyperlinks to other bioinformatics databases | NCBI |
| **NeuroPedia** (18) | 2011 | Chordates | 847 neuropeptides, genomic and taxonomic information, mass spectral libraries (3,401 identified spectra) | www.neuropeptides.nl,<br>Handbook of Biologically Active Peptides, UniProt |
| **NeuroPep 2.0**<br>(19,20) | v1:<br>2015,<br>v2:<br>2024 | Multi | >11,400 neuropeptides with 693 entries from *Crustacea*,(21) 81 families, 924 species, receptor information, predicted and experimental 3D structures, neuropeptide prediction tools (DeepNeuropePred, NeuroPred-PLM) | Uniprot, Neuropedia, PubMed, Europe PMC |
| **DIneR** (22) | 2017 | Insect | 4,782 neuropeptides, 53 families, 425 insect species, physiological functionality, receptor-binding images, family summaries, cladograms, sequence alignments | NCBI |
| **cNPDB**<br>(this work) | 2025 | Crustacean | Comprehensive for *Crustacea*: 1364 neuropeptides, 55 families, 29 species, physiological functionality properties, MSI images, predicted 3D structures | PubMed, Europe PMC, NCBI, NeuroPep |

**Figure 1:**
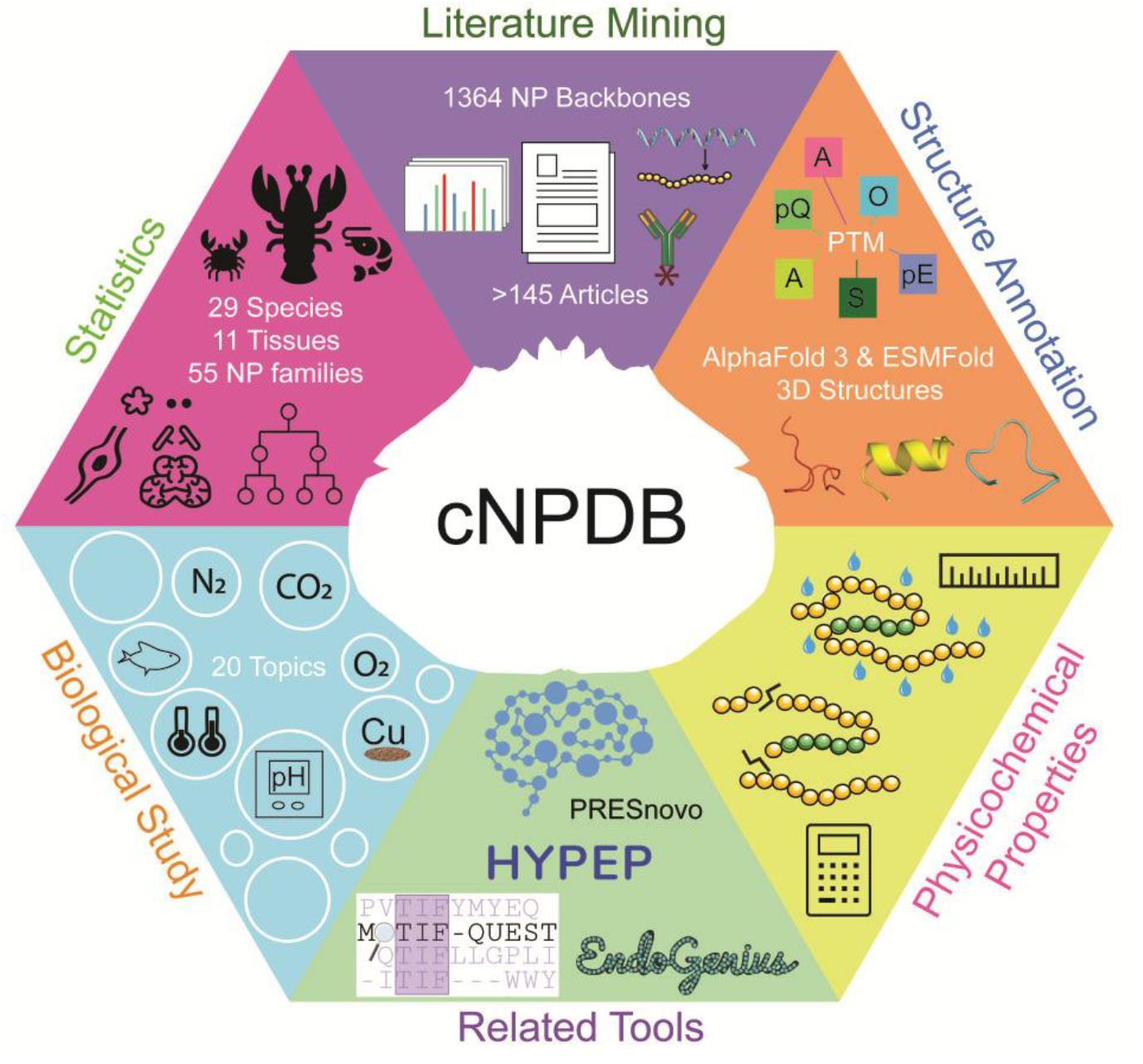
Highlighted content and features in the Crustacean Neuropeptide Database (cNPDB) as the most comprehensive resource for crustacean neuropeptide research with empirical data from literature and built-in tools.

## Materials and Methods

### Software Implementation

The cNPDB web application was implemented using Streamlit (v 1.29.0), a Python-based framework designed for building interactive data-driven interfaces. The source code is publicly available on GitHub (https://github.com/lingjunli-research/CNPDB), and the application is deployed via Streamlit Cloud (https://lilabs.org/cnpdb).

### Data Curation

cNPDB currently contains 1364 crustacean neuropeptides sequences. Of these, 488 entries were sourced from a compiled list in a recently-published review article.(21) The remaining have been reported extensively in the literature, originated initially from bioassays, transcriptomic analysis, mass spectrometry, and other analytical approaches.(26,31,37,41,53-155) Duplicated sequences were removed and reported PTMs (*i*.*e*., N-termini acetylation, oxidation, C-termini amidation, sulfation, N-termini cyclization of glutamine or glutamic acid) were retained in the peptide descriptions. Critical information for each entry, including family classification, organism, and associated neural tissue, were included by cross-referencing with their original articles. Existence level classifications include “predicted”, peptides identified by transcriptome mining exclusively; “*De novo*”, peptides discovered from MS^2^ alone; “MS”, peptides identified solely by precursor mass (MS^1^); and “MSMS”, peptides identified via database searching with MS^1^ and MS2 evidence.

A Python script was developed to automatically query published articles pertaining to each peptide entries against NCBI PubMed, Europe PMC, and NeuroPep. For PubMed, the Entrez package of BioPython (v1.76) was utilized. If flagged when a sequence appeared in titles, abstracts, or full text, PubMed and PMC hits were manually filtered to remove irrelevant references. As an API is unavailable for searching NeuroPep, databases files for eight most well-represented crustacean species (*Callinectes sapidus, Cancer borealis, Carcinus maenas, Homarus americanus, Litopenaeus vannamei, Ocypode ceratophthalma, Orconectes limosus, Procambarus clarkii*) in NeuroPep were downloaded directly on June 12, 2025 and supplemented with manual matches. MS ion images acquired under control conditions were manually selected and reposted with permission from previous publications. (61,64,76,77,79,113,127,134,147,155)

### Physicochemical Property and Structural Predictions

All physicochemical properties for each entry were computed using the Bio.SeqUtils.ProtParam module from BioPython (v1.85), unless specified otherwise.(156)

i. Monoisotopic Mass: [M+H]^+^ *m/z* value were calculated at (http://db.systemsbiology.net/proteomicsToolkit/FragIonServlet.html).
ii. Length: via *len()* function; referring to the number of residues per sequence.
iii. GRAVY score: via *gravy()* function; sum of hydropathicity values divided by peptide length.
iv. Instability index: via *instability_index()* function.
v. Isoelectricpoint (pI): via *isoelectric_point()* function.
vi. Net charge (pH 7.0): via *charge_at_pH(7*.*0)* function.
vii. %Hydrophobic Residues: number of hydrophobic residues divided by total sequence length.
viii. Aliphatic index: a measure of the relative volume occupied by aliphatic side chains in a peptide, following Ikai’s formula.(157)
ix. Boman index: total free energy of amino acids via *boman_scale*.*get()* function divided by peptide length.(158); higher values suggest stronger peptide-protein interactions.

Neuropeptide structures were predicted via AlphaFold 3 and ESMFold using unmodified sequences. AlphaFold 3 predictions were retrieved via the DeepMind (Google) online server, (alphafoldserver.com; accessed June 22, 2025) with the highest scoring structure reported in cNPDB.(159) AlphaFold 3 requires peptide sequences to contain at least four amino acids; therefore, peptides LPA, EDV, and AVG (with cNPDB IDs 742, 746, and 784, respectively) could not be used for 3D structure prediction. Neuropeptides predictions from ESMFold were retrieved from the ESM Metagenomic Atlas (Meta AI) via API (esmatlas.com; accessed June 22, 2025).(160)

### Database structure

#### Home page

The *Home* page provides a broad overview of cNPDB, highlighting its key features, available resources, and how users can stay informed about cNPDB updates.

#### Search Engine page

The search engine allows users to filter neuropeptides by using slider bars and multi-selection boxes, covering physicochemical ranges, biological context or experimental approaches (**Figure 2A**). If no filters are applied, the entire database is displayed for unrestricted browsing. Matching results appear as peptide cards displaying sequence, family, and species (**Figure 2B**). Every card includes a checkbox for selection, with an optional button for batch viewing and download. Once peptide cards are selected, users gain access to detailed information, including physicochemical profiles, literature references, predicted 3D structures (AlphaFold 3, ESMFold) with interactive visualization, and mass spectrometry imaging (MSI) data where available (**Figure 2C**). 3D structure files are downloadable for further analysis using molecular modeling tools and allows a side-by-side prediction comparison between these two models. Users can download the complete search results in multiple formats, including taxonomic, physicochemical and functional profiles (.xlsx), peptide sequences (.FASTA), or a ZIP archive containing all associated 3D structures and MSI files. When no search criteria are applied, users also have the option to download the entire cNPDB dataset.

**Figure 2:**
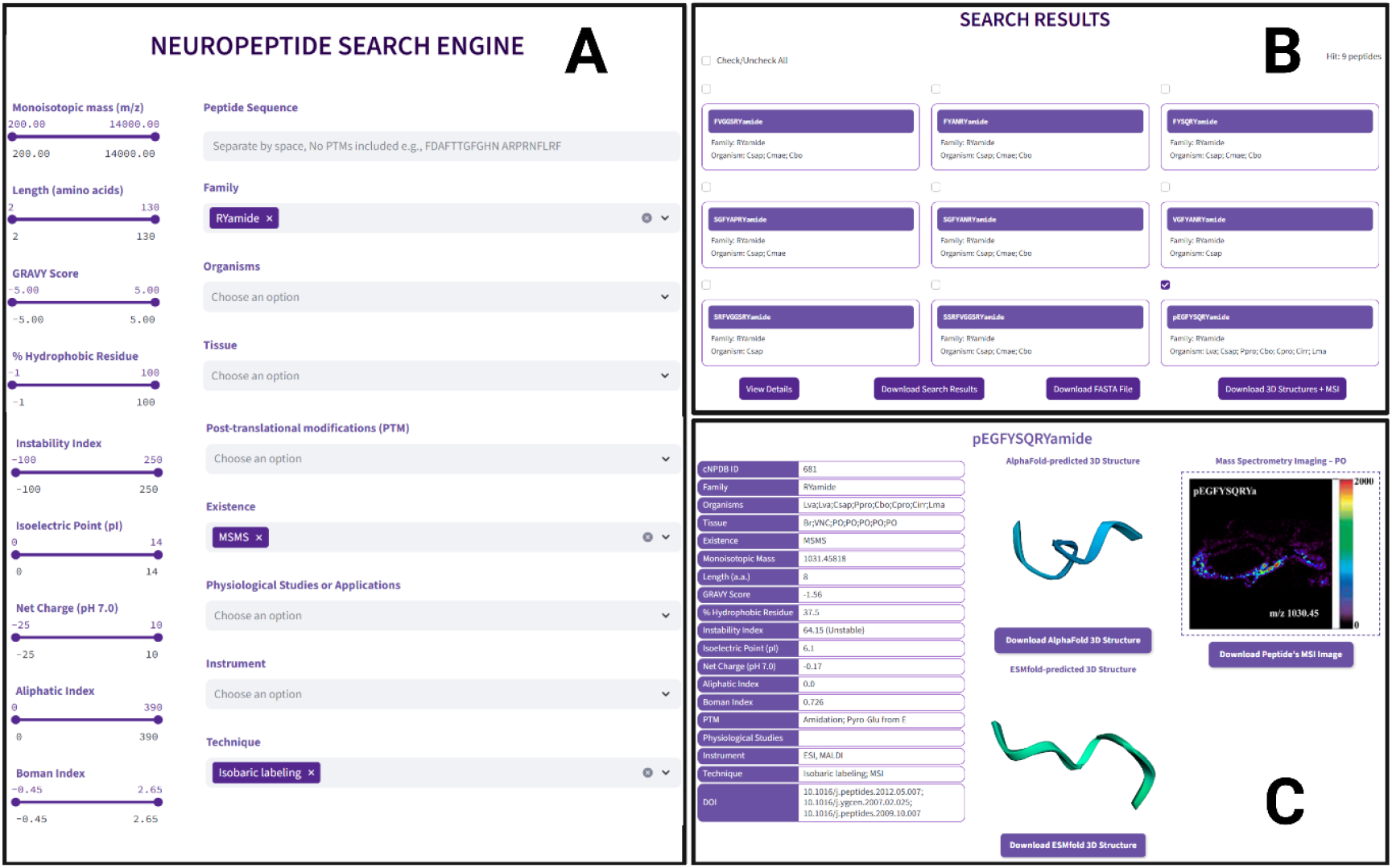
The cNPDB search engine interface. **A)** Representative search using filters “RYamide” family, “MSMS” existence, and “Isobaric labeling” technique. **B)**. Nine matching hits are displayed. **C.)** Representative information page for pEGFYSQRYamide showing its physicochemical properties, predicted 3D structures, and MSI distribution in the *Callinectes sapidus* pericardial organ.

#### Tools page

The peptide property calculator and sequence alignment tools enhance cNPDB’s functionality beyond static browsing. Users can input any peptide to retrieve physicochemical profiles or perform sequence alignments, either against a chosen peptide or the entire cNPDB database, with customizable or well-defined default parameters. Alignment results are presented in a table organized by score or percent identity with detailed graphics and are downloadable for offline analysis. These tools address the challenge of high sequence similarity in neuropeptidome studies,(161,162) helping researchers quickly identify related peptides in their samples.

#### Other pages

The *Related resources* page provides brief descriptions and direct links to other publicly available neuropeptide databases and external tools commonly used in neuropeptide research, mass spectrometry analysis, structural prediction, and homology search.(159-166) The *Statistics* page offers visual charts illustrating the number of cNPDB neuropeptides by species, family, physicochemical properties, biological applications, and experimental techniques, giving researchers a quick understanding of the database’s scope and coverage. The *Tutorial* page offers step-by-step video guides on cNPDB main features, including search engine, filtering criteria and result interpretation, tools, and general website navigation. The *Glossary* page defines species, peptide family, and tissue abbreviations, as well as key concepts related to the calculator and alignment tools. The *FAQs* page addresses common questions and troubleshooting tips for search, downloads, and tool use. The *Contact Us* page provides contact information of the cNPDB development team and highlights key neuropeptidomics publications from our lab. The *Submission* page offers a form for questions, suggestions, technical issues, or new dataset contributions. All submissions are reviewed and addressed promptly. Collectively, these elements contribute to a well-rounded and user-friendly platform, positioning cNPDB as not only a specialized neuropeptide resource, but a practical tool for broader peptide-related research.

## Results and Discussion

### Features of cNPDB neuropeptides

To assess structural stability across cNPDB, we analyzed the distribution of key physicochemical properties. The instability index showed most peptides span from moderately unstable to fairly stable (**Figure 3A**), while the aliphatic index suggested moderate thermostability of most neuropeptides within the database (**Figure 3B**). The grand average of hydropathicity (GRAVY) scores indicated a general tendency toward hydrophilicity (**Figure 3C**), a potentially favorable trait for drug development. The Boman index implied low protein-binding potential and greater specificity (**Figure 3D**). Additional analyses revealed broad biochemical diversity, with most peptides between 5 to 15 amino acids long (**Figure 3E**) and precursor *m/z* values between 400 to 1200 (**Figure 3F**), broadly covering the typical ranges reported in previous studies.(102,163) The net charge at pH 7.0 displayed a nearly-symmetrical distribution around zero, reflecting the electrostatic diversity of peptides that may influence their interactions with G protein-coupled receptors (GPCRs) (**Figure 3G**), while isoelectric points (pI) spanned from acidic to basic ranges (**Figure 3H**), reflecting diverse functional roles across species and tissues. Altogether, these features underscore the structural and biochemical diversity of the cNPDB dataset.

**Figure 3:**
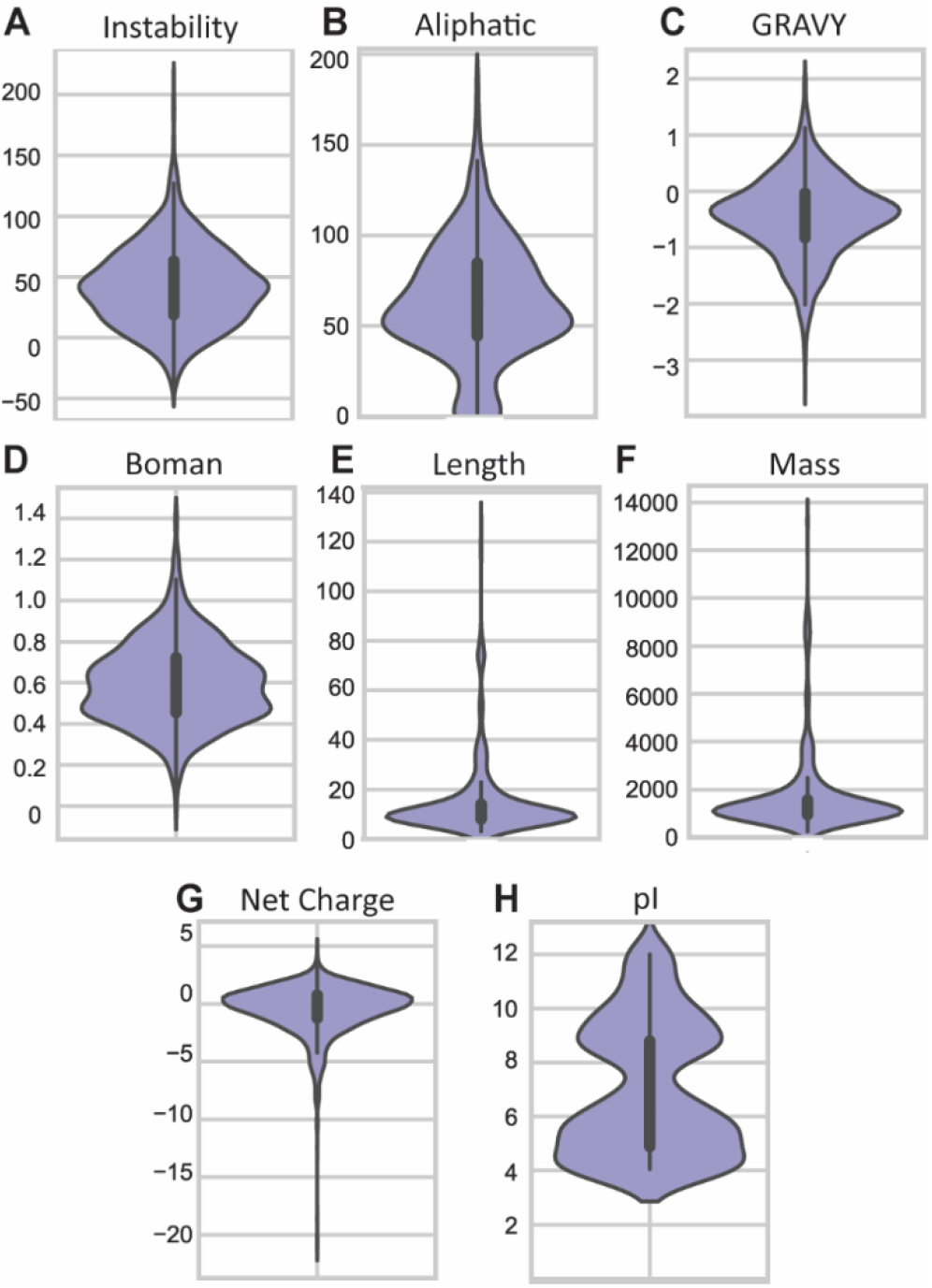
Violin plots showing the distribution of physicochemical properties of neuropeptides in cNPDB, including **A)** instability index, **B)** aliphatic index, **C)** hydrophobicity (GRAVY score), **D)** Boman index, **E)** length, **F)** monoisotopic mass, **G)** net charge at pH 7.0, and **H)** isoelectric point (pI). These properties provide insight into the structural stability and biochemical diversity of crustacean neuropeptides curated in the database.

### Evaluation of Neuropeptide structures

Due to the scarcity of experimentally resolved neuropeptide structures, particularly none in decapods, we aimed to generate putative structural predictions for crustacean neuropeptides. This gap is partly due to their low *in vivo* concentrations and structural complexity.(167) It must be noted that, without experimental references, AlphaFold 3 predictions could not be benchmarked, though reported predicted local distance difference test (pLDDT) scores were generally high and consistent with prior findings.(19) Additionally, we also displayed ESMFold-predicted structures to provide model comparison capabilities. Observed structural patterns included hydrophilic-to-hydrophobic transitions corresponding to disordered-to-helical regions (**Figure 4A**); helical regions flanked on both sides by disordered segments (**Figure 4B**); well-organized, fully helical structures (**Figure 4C-D**); putative cyclic peptide structures (**Figure 4E**); and fully disordered with no apparent secondary structure (**Figure 4F**). In general, lower confidence and more flexibility was observed toward the termini of each peptide (**Figure 4G**), a feature previously linked to enhanced solubility in human neuropeptides.(168) As more prohormones and receptor sequences are identified through genome sequencing, pinpointing the active regions of peptides and their structures will become increasingly important.

**Figure 4:**
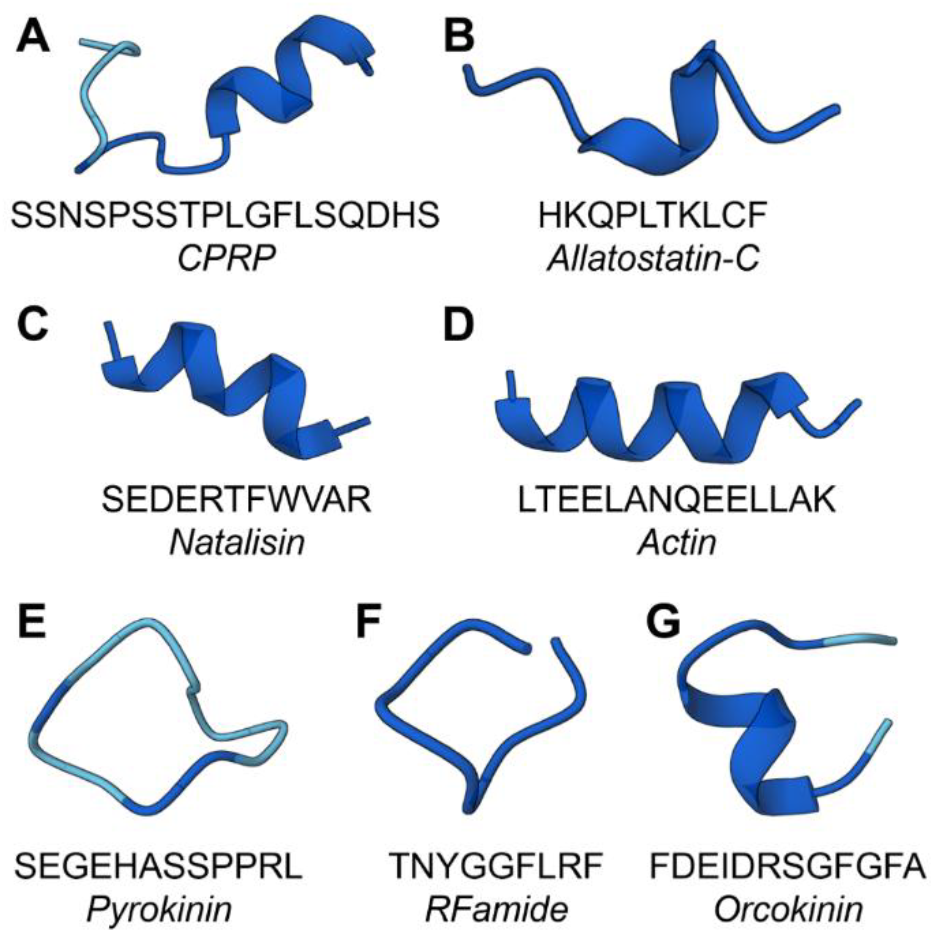
Structures of representative peptides and their corresponding families from as found in cNPDB. **A)** SSNSPSSTPLGFLSQDHS, a crustacean cardioactive peptide precursor related peptide. **B)** HKQPLTKLCF, an allatostatin-C peptide. **C)** SEDERTFWVAR, a natalisin peptide. **D)** LTEELANQEELLAK, an actin peptide. **E)** SEGEHASSPPRL, a pyrokinin peptide. **F)** TNYGGFLRF, an RFamide peptide. **G)** FDEIDRSGFGFA, an orcokinin peptide.

### Usage scenarios

Recognizing that one of the primary rationales for crustacean usage in neuropeptides research is driven by biological applications, we assigned relevant keywords to each peptide based on their source publications. These span fundamental neurobiology studies in cardiac function,(169-171) and immune response,(172,173) as well as behavioral applications including feeding(1,31,169) and circadian rhythm, and others.(174-176) Environmental studies are perhaps the most abundant, covering hypoxia,(37,61,177) salinity levels,(178-180) temperature,(67,180) hypercapnia,(88,177) and copper toxicity(181). We also include research on development and reproduction, including developmental stages,(119,182,183) sexual dimorphism,(184) and vitellogenesis.(56,185) By providing clickable keywords within the database in relation to these applications, methodologies, and instrumentation, even those new to the field of neuropeptidomics can browse a well-documented resource for experimental design. Beyond neuropeptides, crustaceans are widely used for genomic studies,(186-189) neurotransmitter assessment,(190-192) motor pattern probing,(193-195) and transcriptomics.(196,197) Thus, by integrating these topical components into cNPDB, we enable researchers working with crustacea in other mediums to explore potential neuropeptides present within their own work. Our database also displays previously published mass spectrometry imaging mapping neuropeptides within crustacean neural tissues, which offers spatial insights into physiological conditions, including potential co-modulatory activity.(198) Collectively, cNPDB serves as the first comprehensive, dedicated, empirical resource for crustacean neuropeptide research.

## Conclusion

Despite significant advances from crustacean research, the absence of a comprehensive, centralized crustacean neuropeptide database has hindered research progress, as existing resources are often incomplete and lack crucial chemical and biological data. cNPDB fills this gap by providing a freely accessible, user-friendly platform with robust search capabilities, integrated analytical tools, and a wealth of complementary resources, empowering researchers to efficiently explore, analyze, and visualize neuropeptide data. As crustacean genomic resources grow, cNPDB will evolve accordingly to include newly identified neuropeptides and receptors, along with complex PTMs, such as D-amino acid isomerization and glycosylation, whose roles in modulating neuropeptide function are a subject of growing research.(39,41,102) Future updates will also feature enhanced BLAST capabilities for refined homology searches, further supporting evolutionary and functional studies.

## Acknowledgements

Figure 2 was generated with BioRender

## Author Contributions Statement

Conceptualization: V.N.H.T, T.U.D, L.F., and L.L. Bioinformatics: V.N.H.T, T.U.D, and L.F. Data formatting: K.T. and M.B. Funding acquisition: L.L. Supervision: L.L. Writing-original draft: V.N.H.T, T.U.D, L.F., and L.L. Writing-review and editing: all authors.

## References

1. DeLaney, K., Hu, M., Hellenbrand, T., Dickinson, P.S., Nusbaum, M.P. and Li, L. (2021) Mass Spectrometry Quantification, Localization, and Discovery of Feeding-Related Neuropeptides in Cancer borealis. ACS Chemical Neuroscience, 12, 782–798.

2. Wu, W., Ma, M., Ibarra, A.E., Lu, G., Bakshi, V.P. and Li, L. (2023) Global Neuropeptidome Profiling in Response to Predator Stress in Rat: Implications for Post-Traumatic Stress Disorder. Journal of the American Society for Mass Spectrometry, 34, 1549–1558.

3. Lu, G., Ma, F., Wei, P., Ma, M., Tran, V.N.H., Baldo, B.A. and Li, L. (2025) Cocaine-Induced Remodeling of the Rat Brain Peptidome: Quantitative Mass Spectrometry Reveals Anatomically Specific Patterns of Cocaine-Regulated Peptide Changes. ACS Chemical Neuroscience, 16, 128–140.

4. Wang, X., Wang, Q., Zhao, M., Xu, Y., Fu, B., Zhang, L., Wu, S., Yang, D. and Jia, C. (2023) Cold Exposure–induced Alterations in the Brain Peptidome and Gut Microbiome Are Linked to Energy Homeostasis in Mice. Molecular & Cellular Proteomics, 22, 100525.

5. Bao, C., Yang, Y., Huang, H. and Ye, H. (2018) Inhibitory Role of the Mud Crab Short Neuropeptide F in Vitellogenesis and Oocyte Maturation via Autocrine/Paracrine Signaling. Frontiers in Endocrinology, 9, 390.

6. Yeo, X.Y., Cunliffe, G., Ho, R.C., Lee, S.S. and Jung, S. (2022) Potentials of Neuropeptides as Therapeutic Agents for Neurological Diseases. 10, 343.

7. McGonigle, P. (2012) Peptide therapeutics for CNS indications. Biochemical Pharmacology, 83, 559–566.

8. Fosgerau, K. and Hoffmann, T. (2015) Peptide therapeutics: current status and future directions. Drug Discovery Today, 20, 122–128.

9. Brothers, S.P. and Wahlestedt, C. (2010) Therapeutic potential of neuropeptide Y (NPY) receptor ligands. EMBO Molecular Medicine, 2, 429-439-439.

10. Edvinsson, L. (2018) The CGRP Pathway in Migraine as a Viable Target for Therapies. 58, 33–47.

11. Chen, X.-Y., Du, Y.-F. and Chen, L. (2019) Neuropeptides Exert Neuroprotective Effects in Alzheimer’s Disease. Volume 11 - 2018.

12. Enman, N.M., Sabban, E.L., McGonigle, P. and Van Bockstaele, E.J. (2015) Targeting the neuropeptide Y system in stress-related psychiatric disorders. Neurobiology of Stress, 1, 33–43.

13. Ibarra, A.E., Wu, W.X., Zhang, H.R. and Li, L.J. (2025) Quantitative neuropeptide analysis by mass spectrometry: advancing methodologies for biological discovery. Rsc Chem Biol.

14. Hooper, S.L. and DiCaprio, R.A. (2004) Crustacean Motor Pattern Generator Networks. Neurosignals, 13, 50–69.

15. Christie, A.E., Stemmler, E.A. and Dickinson, P.S. (2010) Crustacean neuropeptides. Cellular and Molecular Life Sciences, 67, 4135–4169.

16. Fields, L., Dang, T.C., Tran, V.N.H., Ibarra, A.E. and Li, L. (2025) Decoding Neuropeptide Complexity: Advancing Neurobiological Insights from Invertebrates to Vertebrates through Evolutionary Perspectives. ACS Chemical Neuroscience, 16, 1662–1679.

17. Burbach, J.P. (2010) Neuropeptides from concept to online database www.neuropeptides.nl. Eur J Pharmacol, 626, 27–48.

18. Kim, Y., Bark, S., Hook, V. and Bandeira, N. (2011) NeuroPedia: neuropeptide database and spectral library. Bioinformatics, 27, 2772–2773.

19. Wang, M., Wang, L., Xu, W., Chu, Z., Wang, H., Lu, J., Xue, Z. and Wang, Y. (2024) NeuroPep 2.0: An Updated Database Dedicated to Neuropeptide and Its Receptor Annotations. J Mol Biol, 436, 168416.

20. Wang, Y., Wang, M., Yin, S., Jang, R., Wang, J., Xue, Z. and Xu, T. (2015) NeuroPep: a comprehensive resource of neuropeptides. Database (Oxford), 2015, bav038.

21. Fields, L., Dang, T.C., Tran, V.N.H., Ibarra, A.E. and Li, L. (2025) Decoding Neuropeptide Complexity: Advancing Neurobiological Insights from Invertebrates to Vertebrates through Evolutionary Perspectives. ACS Chemical Neuroscience.

22. Yeoh, J.G.C., Pandit, A.A., Zandawala, M., Nässel, D.R., Davies, S.A. and Dow, J.A.T. (2017) DINeR: Database for Insect Neuropeptide Research. Insect Biochem Molec, 86, 9–19.

23. Wittmann, A.C., Benrabaa, S.A.M., Lopez-Ceron, D.A., Chang, E.S. and Mykles, D.L. (2018) Effects of temperature on survival, moulting, and expression of neuropeptide and mTOR signalling genes in juvenile Dungeness crab (Metacarcinus magister). J Exp Biol, 221.

24. Christie, A.E., Pascual, M.G. and Yu, A. (2018) Peptidergic signaling in the tadpole shrimp Triops newberryi: A potential model for investigating the roles played by peptide paracrines/hormones in adaptation to environmental change. Mar Genomics, 39, 45–63.

25. Powell, D.J., Owens, E., Bergsund, M.M., Cooper, M., Newstein, P., Berner, E., Janmohamed, R. and Dickinson, P.S. (2023) The role of feedback and modulation in determining temperature resiliency in the lobster cardiac nervous system. Front Neurosci, 17, 1113843.

26. Liu, Y., Buchberger, A.R., DeLaney, K., Li, Z. and Li, L. (2019) Multifaceted Mass Spectrometric Investigation of Neuropeptide Changes in Atlantic Blue Crab, Callinectes sapidus, in Response to Low pH Stress. Journal of Proteome Research, 18, 2759–2770.

27. Jayasankar, V., Tomy, S. and Wilder, M.N. (2020) Insights on Molecular Mechanisms of Ovarian Development in Decapod Crustacea: Focus on Vitellogenesis-Stimulating Factors and Pathways. Front Endocrinol (Lausanne), 11, 577925.

28. Nguyen, T.V., Rotllant, G.E., Cummins, S.F., Elizur, A. and Ventura, T. (2018) Insights Into Sexual Maturation and Reproduction in the Norway Lobster (Nephrops norvegicus) via in silico Prediction and Characterization of Neuropeptides and G Protein-coupled Receptors. Front Endocrinol (Lausanne), 9, 430.

29. Glendinning, S., Fitzgibbon, Q.P., Smith, G.G. and Ventura, T. (2023) Unravelling the neuropeptidome of the ornate spiny lobster Panulirus ornatus: A focus on peptide hormones and their processing enzymes expressed in the reproductive tissues. Gen Comp Endocrinol, 332, 114183.

30. Zhang, Y., DeLaney, K., Hui, L., Wang, J., Sturm, R.M. and Li, L. (2018) A Multifaceted Mass Spectrometric Method to Probe Feeding Related Neuropeptide Changes in Callinectes sapidus and Carcinus maenas. J Am Soc Mass Spectrom, 29, 948–960.

31. DeLaney, K., Hu, M., Wu, W., Nusbaum, M.P. and Li, L. (2022) Mass spectrometry profiling and quantitation of changes in circulating hormones secreted over time in Cancer borealis hemolymph due to feeding behavior. Analytical and Bioanalytical Chemistry, 414, 533–543.

32. Wei, Y., Lin, D., Xu, Z., Gao, X., Zeng, C. and Ye, H. (2020) A Possible Role of Crustacean Cardioactive Peptide in Regulating Immune Response in Hepatopancreas of Mud Crab. Front Immunol, 11, 711.

33. Xu, Z., Wei, Y., Wang, G. and Ye, H. (2021) B-type allatostatin regulates immune response of hemocytes in mud crab Scylla paramamosain. Dev Comp Immunol, 120, 104050.

34. Zhai, B., Li, X., Lin, C., Yan, P., Zhao, Q. and Li, E. (2021) Proteomic analysis of hemocyte reveals the immune regulatory mechanisms after the injection of corticosteroid-releasing hormone in mud crab Scylla Paramamosain. J Proteomics, 242, 104238.

35. Wei, Y., Xu, Z., Hao, S., Guo, S., Huang, H. and Ye, H. (2022) Immunomodulatory role of crustacean cardioactive peptide in the mud crab Scylla paramamosain. Fish Shellfish Immunol, 121, 142–151.

36. Buchberger, A.R., DeLaney, K., Liu, Y., Vu, N.Q., Helfenbein, K. and Li, L. (2020) Mass Spectrometric Profiling of Neuropeptides in Callinectes sapidus during Hypoxia Stress. ACS Chem Neurosci, 11, 3097–3106.

37. Buchberger, A.R., Sauer, C.S., Vu, N.Q., DeLaney, K. and Li, L. (2020) Temporal Study of the Perturbation of Crustacean Neuropeptides Due to Severe Hypoxia Using 4-Plex Reductive Dimethylation. Journal of Proteome Research, 19, 1548–1555.

38. Christie, A.E., Miller, A., Fernandez, R., Dickinson, E.S., Jordan, A., Kohn, J., Youn, M.C. and Dickinson, P.S. (2018) Non-amidated and amidated members of the C-type allatostatin (AST-C) family are differentially distributed in the stomatogastric nervous system of the American lobster, Homarus americanus. Invert Neurosci, 18, 2.

39. Phetsanthad, A., Roycroft, C. and Li, L. (2023) Enrichment and fragmentation approaches for enhanced detection and characterization of endogenous glycosylated neuropeptides. Proteomics, 23, e2100375.

40. Ollivaux, C., Gallois, D., Amiche, M., Boscameric, M. and Soyez, D. (2009) Molecular and cellular specificity of post-translational aminoacyl isomerization in the crustacean hyperglycaemic hormone family. FEBS J, 276, 4790–4802.

41. Tran, V.N.H., Lu, G., Duong, T., Ibarra, A.E., Wu, F., Beaver, M. and Li, L. (2025) HILIC-Enabled Mass Spectrometric Discovery of Novel Endogenous and Glycosylated Neuropeptides in the American Lobster Nervous System. bioRxiv.

42. Muscato, A.J., Powell, D.J., Bulhan, W., Mackenzie, E.S., Pupo, A., Rolph, M., Christie, A.E. and Dickinson, P.S. (2022) Structural variation between neuropeptide isoforms affects function in the lobster cardiac system. Gen Comp Endocrinol, 327, 114065.

43. Oleisky, E.R., Stanhope, M.E., Hull, J.J. and Dickinson, P.S. (2022) Isoforms of the neuropeptide myosuppressin differentially modulate the cardiac neuromuscular system of the American lobster, Homarus americanus. J Neurophysiol, 127, 702–713.

44. Serrano, L., Blanvillain, G., Soyez, D., Charmantier, G., Grousset, E., Aujoulat, F. and Spanings-Pierrot, C. (2003) Putative involvement of crustacean hyperglycemic hormone isoforms in the neuroendocrine mediation of osmoregulation in the crayfish Astacus leptodactylus. J Exp Biol, 206, 979–988.

45. Tom, M., Manfrin, C., Mosco, A., Gerdol, M., De Moro, G., Pallavicini, A. and Giulianini, P.G. (2014) Different transcription regulation routes are exerted by L- and D-amino acid enantiomers of peptide hormones. J Exp Biol, 217, 4337–4346.

46. Soyez, D., Van Herp, F., Rossier, J., Le Caer, J.P., Tensen, C.P. and Lafont, R. (1994) Evidence for a conformational polymorphism of invertebrate neurohormones. D-amino acid residue in crustacean hyperglycemic peptides. J Biol Chem, 269, 18295–18298.

47. Gnanabharathi, B., Fahoum, S.H. and Blitz, D.M. (2024) Neuropeptide Modulation Enables Biphasic Internetwork Coordination via a Dual-Network Neuron. eNeuro, 11.

48. Fahoum, S.H. and Blitz, D.M. (2024) Neuropeptide modulation of bidirectional internetwork synapses. J Neurophysiol, 132, 184–205.

49. Cronin, E.M., Schneider, A.C., Nadim, F. and Bucher, D. (2024) Modulation by Neuropeptides with Overlapping Targets Results in Functional Overlap in Oscillatory Circuit Activation. J Neurosci, 44.

50. Hyde, C.J., Nguyen, T., Fitzgibbon, Q.P., Elizur, A., Smith, G.G. and Ventura, T. (2020) Neural remodelling in spiny lobster larvae is characterized by broad neuropeptide suppression. Gen Comp Endocrinol, 294, 113496.

51. Chen, X., Wang, J., Hou, X., Yue, W., Huang, S. and Wang, C. (2019) Tissue expression profiles unveil the gene interaction of hepatopancreas, eyestalk, and ovary in the precocious female Chinese mitten crab, Eriocheir sinensis. BMC Genet, 20, 12.

52. Dickinson, P.S., Hull, J.J., Miller, A., Oleisky, E.R. and Christie, A.E. (2019) To what extent may peptide receptor gene diversity/complement contribute to functional flexibility in a simple pattern-generating neural network? Comp Biochem Physiol Part D Genomics Proteomics, 30, 262–282.

53. Aguilar, M.B., Soyez, D., Falchetto, R., Arnott, D., Shabanowitz, J., Hunt, D.F. and Huberman, A. (1995) Amino acid sequence of the minor isomorph of the crustacean hyperglycemic hormone (CHH-II) of the Mexican crayfish Procambarus bouvieri (Ortmann): Presence of a d-amino acid. Peptides, 16, 1375–1383.

54. Alexander, J.L., Oliphant, A., Wilcockson, D.C., Audsley, N., Down, R.E., Lafont, R. and Webster, S.G. (2018) Functional Characterization and Signaling Systems of Corazonin and Red Pigment Concentrating Hormone in the Green Shore Crab, Carcinus maenas. Frontiers in Neuroscience, 11, 752.

55. Audsley, N. and Weaver, R.J. (2003) Identification of neuropeptides from brains of larval Manduca sexta and Lacanobia oleracea using MALDI-TOF mass spectrometry and post-source decay. Peptides, 24, 1465–1474.

56. Bao, C., Yang, Y., Huang, H. and Ye, H. (2015) Neuropeptides in the cerebral ganglia of the mud crab, Scylla paramamosain: transcriptomic analysis and expression profiles during vitellogenesis. Sci Rep, 5, 17055.

57. Behrens, H.L., Chen, R. and Li, L. (2008) Combining Microdialysis, NanoLC-MS, and MALDI-TOF/TOF To Detect Neuropeptides Secreted in the Crab, Cancer borealis. Analytical Chemistry, 80, 6949–6958.

58. Billimoria, C.P., Li, L. and Marder, E. (2005) Profiling of neuropeptides released at the stomatogastric ganglion of the crab, Cancer borealis with mass spectrometry. Journal of Neurochemistry, 95, 191–199.

59. Bungart, D., Hilbich, C., Dircksen, H. and Keller, R. (1995) Occurrence of analogues of the myotropic neuropeptide orcokinin in the shore crab, Carcinus maenas: Evidence for a novel neuropeptide family. Peptides, 16, 67–72.

60. Cao, Q., Yu, Q., Liu, Y., Chen, Z. and Li, L. (2020) Signature-Ion-Triggered Mass Spectrometry Approach Enabled Discovery of N- and O-Linked Glycosylated Neuropeptides in the Crustacean Nervous System. Journal of Proteome Research, 19, 634–643.

61. Buchberger, A.R., Vu, N.Q., Johnson, J., DeLaney, K. and Li, L. (2020) A Simple and Effective Sample Preparation Strategy for MALDI-MS Imaging of Neuropeptide Changes in the Crustacean Brain Due to Hypoxia and Hypercapnia Stress. Journal of the American Society for Mass Spectrometry, 31, 1058–1065.

62. Cape, S.S., Rehm, K.J., Ma, M., Marder, E. and Li, L. (2008) Mass spectral comparison of the neuropeptide complement of the stomatogastric ganglion and brain in the adult and embryonic lobster, Homarus americanus. Journal of Neurochemistry, 105, 690–702.

63. Chen, R., Hui, L., Cape, S.S., Wang, J. and Li, L. (2009) Comparative Neuropeptidomic Analysis of Food Intake via a Multifaceted Mass Spectrometric Approach. ACS Chemical Neuroscience, 1, 204–214.

64. Chen, R., Hui, L., Sturm, R.M. and Li, L. (2009) Three dimensional mapping of neuropeptides and lipids in crustacean brain by mass spectral imaging. Journal of the American Society for Mass Spectrometry, 20, 1068–1077.

65. Chen, R., Ma, M., Hui, L., Zhang, J. and Li, L. (2009) Measurement of neuropeptides in crustacean hemolymph via MALDI mass spectrometry. Journal of the American Society for Mass Spectrometry, 20, 708–718.

66. Chen, R., Ouyang, C., Xiao, M. and Li, L. (2014) In situ identification and mapping of neuropeptides from the stomatogastric nervous system of Cancer borealis. Rapid communications in mass spectrometry: RCM, 28, 2437–2444.

67. Chen, R., Xiao, M., Buchberger, A. and Li, L. (2014) Quantitative Neuropeptidomics Study of the Effects of Temperature Change in the Crab Cancer borealis. Journal of Proteome Research, 13, 5767–5776.

68. Chen, T., Ren, C., Wang, Y., Gao, Y., Wong, N.-K., Zhang, L. and Hu, C. (2016) Crustacean cardioactive peptide (CCAP) of the Pacific white shrimp (Litopenaeus vannamei): Molecular characterization and its potential roles in osmoregulation and freshwater tolerance. Aquaculture, 451, 405–412.

69. Christie, A.E., Cashman, C.R., Stevens, J.S., Smith, C.M., Beale, K.M., Stemmler, E.A., Greenwood, S.J., Towle, D.W. and Dickinson, P.S. (2008) Identification and cardiotropic actions of brain/gut-derived tachykinin-related peptides (TRPs) from the American lobster Homarus americanus. Peptides, 29, 1909–1918.

70. Christie, A.E., Chi, M., Lameyer, T.J., Pascual, M.G., Shea, D.N., Stanhope, M.E., Schulz, D.J. and Dickinson, P.S. (2015) Neuropeptidergic Signaling in the American Lobster Homarus americanus: New Insights from High-Throughput Nucleotide Sequencing. PLOS ONE, 10, e0145964.

71. Christie, A.E., Lundquist, C.T., Nässel, D.R. and Nusbaum, M.P. (1997) Two novel tachykinin-related peptides from the nervous system of the crab Cancer borealis. Journal of Experimental Biology, 200, 2279–2294.

72. Christie, A.E. and Pascual, M.G. (2016) Peptidergic signaling in the crab Cancer borealis: Tapping the power of transcriptomics for neuropeptidome expansion. General and Comparative Endocrinology, 237, 53–67.

73. Christie, A.E., Roncalli, V., Cieslak, M.C., Pascual, M.G., Yu, A., Lameyer, T.J., Stanhope, M.E. and Dickinson, P.S. (2017) Prediction of a neuropeptidome for the eyestalk ganglia of the lobster Homarus americanus using a tissue-specific de novo assembled transcriptome. General and Comparative Endocrinology, 243, 96–119.

74. Christie, A.E., Stevens, J.S., Bowers, M.R., Chapline, M.C., Jensen, D.A., Schegg, K.M., Goldwaser, J., Kwiatkowski, M.A., Pleasant, T.K., Shoenfeld, L. et al. (2010) Identification of a calcitonin-like diuretic hormone that functions as an intrinsic modulator of the American lobster, Homarus americanus, cardiac neuromuscular system. Journal of Experimental Biology, 213, 118–127.

75. Chung, J.S., Wilkinson, M.C. and Webster, S.G. (1998) Amino acid sequences of both isoforms of crustacean hyperglycemic hormone (CHH) and corresponding precursor-related peptide in Cancer pagurus. Regulatory Peptides, 77, 17–24.

76. DeKeyser, S.S., Kutz-Naber, K.K., Schmidt, J.J., Barrett-Wilt, G.A. and Li, L. (2007) Imaging Mass Spectrometry of Neuropeptides in Decapod Crustacean Neuronal Tissues. Journal of Proteome Research, 6, 1782–1791.

77. DeLaney, K., Hu, M.Z., Hellenbrand, T., Dickinson, P.S., Nusbaum, M.P. and Li, L.J. (2021) Mass Spectrometry Quantification, Localization, and Discovery of Feeding-Related Neuropeptides in. Acs Chemical Neuroscience, 12, 782–798.

78. DeLaney, K. and Li, L. (2019) Data Independent Acquisition Mass Spectrometry Method for Improved Neuropeptidomic Coverage in Crustacean Neural Tissue Extracts. Analytical Chemistry, 91, 5150–5158.

79. DeLaney, K. and Li, L. (2020) Neuropeptidomic Profiling and Localization in the Crustacean Cardiac Ganglion Using Mass Spectrometry Imaging with Multiple Platforms. Journal of the American Society for Mass Spectrometry, 31, 2469–2478.

80. Dickinson, P.S., Stemmler, E.A., Barton, E.E., Cashman, C.R., Gardner, N.P., Rus, S., Brennan, H.R., McClintock, T.S. and Christie, A.E. (2009) Molecular, mass spectral, and physiological analyses of orcokinins and orcokinin precursor-related peptides in the lobster Homarus americanus and the crayfish Procambarus clarkii. Peptides, 30, 297–317.

81. Dickinson, P.S., Wiwatpanit, T., Gabranski, E.R., Ackerman, R.J., Stevens, J.S., Cashman, C.R., Stemmler, E.A. and Christie, A.E. (2009) Identification of SYWKQCAFNAVSCFamide: a broadly conserved crustacean C-type allatostatin-like peptide with both neuromodulatory and cardioactive properties. Journal of Experimental Biology, 212, 1140–1152.

82. (1999) Structure, distribution, and biological activity of novel members of the allatostatin family in the crayfish Orconectes limosus. Peptides, 20, 695–712.

83. Duve, H., Johnsen, A.H., Scott, A.G. and Thorpe, A. (2002) Allatostatins of the tiger prawn, Penaeus monodon (Crustacea: Penaeidea). Peptides, 23, 1039–1051.

84. Fort, T.J., Brezina, V. and Miller, M.W. (2007) Regulation of the Crab Heartbeat by FMRFamide-Like Peptides: Multiple Interacting Effects on Center and Periphery. Journal of Neurophysiology, 98, 2887–2902.

85. Fu, Q., Goy, M.F. and Li, L. (2005) Identification of neuropeptides from the decapod crustacean sinus glands using nanoscale liquid chromatography tandem mass spectrometry. Biochemical and Biophysical Research Communications, 337, 765–778.

86. Fu, Q., Kutz, K.K., Schmidt, J.J., Hsu, Y.W.A., Messinger, D.I., Cain, S.D., De La Iglesia, H.O., Christie, A.E. and Li, L. (2005) Hormone complement of the Cancer productus sinus gland and pericardial organ: An anatomical and mass spectrometric investigation. Journal of Comparative Neurology, 493, 607–626.

87. Gu, P.-L., Yu, K.L. and Chan, S.-M. (2000) Molecular characterization of an additional shrimp hyperglycemic hormone: cDNA cloning, gene organization, expression and biological assay of recombinant proteins^1^. FEBS Letters, 472, 122–128.

88. Hu, M., Helfenbein, K., Buchberger, A.R., DeLaney, K., Liu, Y. and Li, L. (2021) Exploring the Sexual Dimorphism of Crustacean Neuropeptide Expression Using Callinectes sapidus as a Model Organism. Journal of Proteome Research, 20, 2739–2750.

89. Huberman, A., Aguilar, M.B., Brew, K., Shabanowitz, J. and Hunt, D.F. (1993) Primary structure of the major isomorph of the crustacean hyperglycemic hormone (CHH-I) from the sinus gland of the Mexican crayfish Procambarus bouvieri (Ortmann): Interspecies comparison. Peptides, 14, 7–16.

90. Hui, L., Cunningham, R., Zhang, Z., Cao, W., Jia, C. and Li, L. (2011) Discovery and characterization of the Crustacean hyperglycemic hormone precursor related peptides (CPRP) and orcokinin neuropeptides in the sinus glands of the blue crab Callinectes sapidus using multiple tandem mass spectrometry techniques. Journal of Proteome Research, 10, 4219–4229.

91. Hui, L., D’Andrea, B.T., Jia, C., Liang, Z., Christie, A.E. and Li, L. (2013) Mass spectrometric characterization of the neuropeptidome of the ghost crab Ocypode ceratophthalma (Brachyura, Ocypodidae). General and Comparative Endocrinology, 184, 22–34.

92. Hui, L., Xiang, F., Zhang, Y. and Li, L. (2012) Mass spectrometric elucidation of the neuropeptidome of a crustacean neuroendocrine organ. Peptides, 36, 230–239.

93. Hui, L., Zhang, Y., Wang, J., Cook, A., Ye, H., Nusbaum, M.P. and Li, L. (2011) Discovery and Functional Study of a Novel Crustacean Tachykinin Neuropeptide. ACS Chemical Neuroscience, 2, 711–722.

94. Huybrechts, J., Nusbaum, M.P., Bosch, L.V., Baggerman, G., Loof, A.D. and Schoofs, L. (2003) Neuropeptidomic analysis of the brain and thoracic ganglion from the Jonah crab, Cancer borealis. Biochemical and Biophysical Research Communications, 308, 535–544.

95. Jia, C., Hui, L., Cao, W., Lietz, C.B., Jiang, X., Chen, R., Catherman, A.D., Thomas, P.M., Ge, Y., Kelleher, N.L. et al. (2012) High-definition De Novo Sequencing of Crustacean Hyperglycemic Hormone (CHH)-family Neuropeptides. Molecular & Cellular Proteomics, 11, 1951–1964.

96. Jia, C., Lietz, C.B., Ye, H., Hui, L., Yu, Q., Yoo, S. and Li, L. (2013) A multi-scale strategy for discovery of novel endogenous neuropeptides in the crustacean nervous system. Journal of Proteomics, 91, 1–12.

97. Jia, C., Yu, Q., Wang, J. and Li, L. (2014) Qualitative and quantitative top-down mass spectral analysis of crustacean hyperglycemic hormones in response to feeding. PROTEOMICS, 14, 1185–1194.

98. Jiang, X., Xiang, F., Jia, C., Buchberger, A.R. and Li, L. (2018) Relative Quantitation of Neuropeptides at Multiple Developmental Stages of the American Lobster Using N, N -Dimethyl Leucine Isobaric Tandem Mass Tags. ACS Chemical Neuroscience, 9, 2054–2063.

99. Li, L., Kelley, W.P., Billimoria, C.P., Christie, A.E., Pulver, S.R., Sweedler, J.V. and Marder, E. (2003) Mass spectrometric investigation of the neuropeptide complement and release in the pericardial organs of the crab, Cancer borealis. Journal of Neurochemistry, 87, 642–656.

100. Liang, Z., M. Schmerberg, C. and Li, L. (2015) Mass spectrometric measurement of neuropeptide secretion in the crab, Cancer borealis, by in vivo microdialysis. Analyst, 140, 3803–3813.

101. Liu, Y., Li, G. and Li, L. (2021) Targeted Top-Down Mass Spectrometry for the Characterization and Tissue-Specific Functional Discovery of Crustacean Hyperglycemic Hormones (CHH) and CHH Precursor-Related Peptides in Response to Low pH Stress. Journal of the American Society for Mass Spectrometry, 32, 1352–1360.

102. Lu, G.Y., Tran, V.N.H., Wu, W.X., Ma, M. and Li, L.J. (2024) Neuropeptidomics of the American Lobster. Journal of Proteome Research, 23, 1757–1767.

103. Ma, M., Bors, E.K., Dickinson, E.S., Kwiatkowski, M.A., Sousa, G.L., Henry, R.P., Smith, C.M., Towle, D.W., Christie, A.E. and Li, L. (2009) Characterization of the Carcinus maenas neuropeptidome by mass spectrometry and functional genomics. General and Comparative Endocrinology, 161, 320–334.

104. Ma, M., Chen, R., Sousa, G.L., Bors, E.K., Kwiatkowski, M.A., Goiney, C.C., Goy, M.F., Christie, A.E. and Li, L. (2008) Mass spectral characterization of peptide transmitters/hormones in the nervous system and neuroendocrine organs of the American lobster Homarus americanus. General and Comparative Endocrinology, 156, 395–409.

105. Ma, M., Gard, A.L., Xiang, F., Wang, J., Davoodian, N., Lenz, P.H., Malecha, S.R., Christie, A.E. and Li, L. (2010) Combining in silico transcriptome mining and biological mass spectrometry for neuropeptide discovery in the Pacific white shrimp Litopenaeus vannamei. Peptides, 31, 27–43.

106. Ma, M., Szabo, T.M., Jia, C., Marder, E. and Li, L. (2009) Mass spectrometric characterization and physiological actions of novel crustacean C-type allatostatins. Peptides, 30, 1660–1668.

107. Ma, M., Wang, J., Chen, R. and Li, L. (2009) Expanding the Crustacean Neuropeptidome Using a Multifaceted Mass Spectrometric Approach. Journal of Proteome Research, 8, 2426–2437.

108. McGrath, L.L., Vollmer, S.V., Kaluziak, S.T. and Ayers, J. (2016) De novo transcriptome assembly for the lobster Homarus americanus and characterization of differential gene expression across nervous system tissues. BMC Genomics, 17, 63.

109. Mercier, A.J., Orchard, I., TeBrugge, V. and Skerrett, M. (1993) Isolation of two FMRFamide-related peptides from crayfish pericardial organs. Peptides, 14, 137–143.

110. Muscato, A.J., Walsh, P., Pong, S., Pupo, A., Gross, R.J., Christie, A.E., Hull, J.J. and Dickinson, P.S. (2021) Does Differential Receptor Distribution Underlie Variable Responses to a Neuropeptide in the Lobster Cardiac System? International Journal of Molecular Sciences, 22, 8703.

111. Nagasawa, H., Yang, W.-J., Shimizu, H., Aida, K., Tsutsumi, H., Terauchi, A. and Sonobe, H. (1996) Isolation and Amino Acid Sequence of a Molt-inhibiting Hormone from the American Crayfish, Procambarus clarkii. Bioscience, Biotechnology, and Biochemistry, 60, 554–556.

112. Nguyen, T.V., Cummins, S.F., Elizur, A. and Ventura, T. (2016) Transcriptomic characterization and curation of candidate neuropeptides regulating reproduction in the eyestalk ganglia of the Australian crayfish,. Scientific Reports, 6.

113. OuYang, C., Chen, B. and Li, L. (2015) High Throughput In Situ DDA Analysis of Neuropeptides by Coupling Novel Multiplex Mass Spectrometric Imaging (MSI) with Gas-Phase Fractionation. Journal of the American Society for Mass Spectrometry, 26, 1992–2001.

114. OuYang, C., Liang, Z. and Li, L. (2015) Mass Spectrometric Analysis of Spatio-Temporal Dynamics of Crustacean Neuropeptides. Biochimica et biophysica acta, 1854, 798–811.

115. Paine, M.R.L., Ellis, S.R., Maloney, D., Heeren, R.M.A. and Verhaert, P.D.E.M. (2018) Digestion-Free Analysis of Peptides from 30-year-old Formalin-Fixed, Paraffin-Embedded Tissue by Mass Spectrometry Imaging. Analytical Chemistry, 90, 9272–9280.

116. Phetsanthad, A., Carr, A.V., Fields, L. and Li, L. (2023) Definitive Screening Designs to Optimize Library-Free DIA-MS Identification and Quantification of Neuropeptides. Journal of Proteome Research.

117. Porras, M.G., De Loof, A., Breuer, M. and Aréchiga, H. (2003) Corazonin promotes tegumentary pigment migration in the crayfish Procambarus clarkii. Peptides, 24, 1581–1589.

118. Rue, M.C.P., Baas-Thomas, N., Iyengar, P.S., Scaria, L.K. and Marder, E. (2022) Localization of chemical synapses and modulatory release sites in the cardiac ganglion of the crab, Cancer borealis. Journal of Comparative Neurology, 530, 2954–2965.

119. Sauer, C.S. and Li, L. (2021) Mass Spectrometric Profiling of Neuropeptides in Response to Copper Toxicity via Isobaric Tagging. Chemical research in toxicology, 10.1021/acs.chemrestox.1020c00521.

120. Schmerberg, C.M. and Li, L. (2013) Function-Driven Discovery of Neuropeptides with Mass Spectrometry-Based Tools. Protein & Peptide Letters, 20, 681–694.

121. Shimma, S. (2022) Mass Spectrometry Imaging. Mass Spectrometry, 11, A0102–A0102.

122. Soyez, D., Le Caer, J.P., Noel, P.Y. and Rossier, J. (1991) Primary structure of two isoforms of the Vitellogenesis Inhibiting Hormone from the Iobster Homarus americanus. Neuropeptides, 20, 25–32.

123. Stemmler, E.A., Barton, E.E., Esonu, O.K., Polasky, D.A., Onderko, L.L., Bergeron, A.B., Christie, A.E. and Dickinson, P.S. (2013) C-terminal methylation of truncated neuropeptides: An enzyme-assisted extraction artifact involving methanol. Peptides, 46, 108–125.

124. Stemmler, E.A., Bruns, E.A., Cashman, C.R., Dickinson, P.S. and Christie, A.E. (2010) Molecular and mass spectral identification of the broadly conserved decapod crustacean neuropeptide pQIRYHQCYFNPISCF: The first PISCF-allatostatin (Manduca sexta- or C-type allatostatin) from a non-insect. General and Comparative Endocrinology, 165, 1–10.

125. Stemmler, E.A., Bruns, E.A., Gardner, N.P., Dickinson, P.S. and Christie, A.E. (2007) Mass spectrometric identification of pEGFYSQRYamide: A crustacean peptide hormone possessing a vertebrate neuropeptide Y (NPY)-like carboxy-terminus. General and Comparative Endocrinology, 152, 1–7.

126. Stemmler, E.A., Peguero, B., Bruns, E.A., Dickinson, P.S. and Christie, A.E. (2007) Identification, physiological actions, and distribution of TPSGFLGMRamide: a novel tachykinin-related peptide from the midgut and stomatogastric nervous system of Cancer crabs. Journal of Neurochemistry, 101, 1351–1366.

127. Sturm, R.M., Greer, T., Chen, R., Hensen, B. and Li, L. (2013) Comparison of NIMS and MALDI platforms for neuropeptide and lipid mass spectrometric imaging in C. borealis brain tissue. Analytical Methods, 5, 1623.

128. Sturm, R.M., Greer, T., Woodards, N., Gemperline, E. and Li, L. (2013) Mass spectrometric evaluation of neuropeptidomic profiles upon heat stabilization treatment of neuroendocrine tissues in crustaceans. Journal of proteome research, 12, 743–752.

129. Suwansa-ard, S., Thongbuakaew, T., Wang, T., Zhao, M., Elizur, A., Hanna, P.J., Sretarugsa, P., Cummins, S.F. and Sobhon, P. (2015) In silico Neuropeptidome of Female Macrobrachium rosenbergii Based on Transcriptome and Peptide Mining of Eyestalk, Central Nervous System and Ovary. PLOS ONE, 10, e0123848.

130. Szabo, T.M., Chen, R., Goeritz, M.L., Maloney, R.T., Tang, L.S., Li, L. and Marder, E. (2011) Distribution and physiological effects of B-type allatostatins (myoinhibitory peptides, MIPs) in the stomatogastric nervous system of the crab Cancer borealis. Journal of Comparative Neurology, 519, 2658–2676.

131. Toullec, J.-Y., Corre, E., Bernay, B., Thorne, M.A.S., Cascella, K., Ollivaux, C., Henry, J. and Clark, M.S. (2013) Transcriptome and Peptidome Characterisation of the Main Neuropeptides and Peptidic Hormones of a Euphausiid: The Ice Krill, Euphausia crystallorophias. PLoS ONE, 8, e71609.

132. Tu, S., Xu, R., Wang, M., Xie, X., Bao, C. and Zhu, D. (2021) Identification and characterization of expression profiles of neuropeptides and their GPCRs in the swimming crab, Portunus trituberculatus. PeerJ, 9, e12179.

133. Ventura, T., Cummins, S.F., Fitzgibbon, Q., Battaglene, S. and Elizur, A. (2014) Analysis of the Central Nervous System Transcriptome of the Eastern Rock Lobster Sagmariasus verreauxi Reveals Its Putative Neuropeptidome. PLoS ONE, 9, e97323.

134. Vu, N.Q., Buchberger, A.R., Johnson, J. and Li, L. (2021) Complementary neuropeptide detection in crustacean brain by mass spectrometry imaging using formalin and alternative aqueous tissue washes. Analytical and Bioanalytical Chemistry, 413, 2665–2673.

135. Wang, J., Ma, M., Chen, R. and Li, L. (2008) Enhanced Neuropeptide Profiling via Capillary Electrophoresis Off-Line Coupled with MALDI FTMS. Analytical Chemistry, 80, 6168–6177.

136. Wang, J., Ye, H., Zhang, Z., Xiang, F., Girdaukas, G. and Li, L. (2011) Advancing Matrix-Assisted Laser Desorption/Ionization-Mass Spectrometric Imaging for Capillary Electrophoresis Analysis of Peptides. Analytical Chemistry, 83, 3462–3469.

137. Wang, L., Huang, C., Wang, M.X., Xue, Z.D. and Wang, Y. (2023) NeuroPred-PLM: an interpretable and robust model for neuropeptide prediction by protein language model. Brief Bioinform,

138. Wei, Y., Lin, D., Xu, Z., Gao, X., Zeng, C. and Ye, H. (2020) A Possible Role of Crustacean Cardioactive Peptide in Regulating Immune Response in Hepatopancreas of Mud Crab. Frontiers in Immunology, 11, 711.

139. Wilcockson, D.C., Zhang, L., Hastings, M.H., Kyriacou, C.P. and Webster, S.G. (2011) A novel form of pigment-dispersing hormone in the central nervous system of the intertidal marine isopod, Eurydice pulchra (leach). Journal of Comparative Neurology, 519, 562–575.

140. Xiang, F., Ye, H., Chen, R., Fu, Q. and Li, L. (2010) N, N -Dimethyl Leucines as Novel Isobaric Tandem Mass Tags for Quantitative Proteomics and Peptidomics. Analytical Chemistry, 82, 2817–2825.

141. Yasuda, A., Yasuda, Y., Fujita, T. and Naya, Y. (1994) Characterization of Crustacean Hyperglycemic Hormone from the Crayfish (Procambarus clarkii): Multiplicity of Molecular Forms by Stereoinversion and Diverse Functions. General and Comparative Endocrinology, 95, 387–398.

142. Yasuda, A., Yasuda-Kamatani, Y., Nozaki, M. and Nakajima, T. (2004) Identification of GYRKPPFNGSIFamide (crustacean-SIFamide) in the crayfish Procambarus clarkii by topological mass spectrometry analysis. General and Comparative Endocrinology, 135, 391–400.

143. Yasuda-Kamatani, Y. and Yasuda, A. (2000) Identification of Orcokinin Gene-Related Peptides in the Brain of the Crayfish Procambarus clarkii by the Combination of MALDI-TOF and On-Line Capillary HPLC/Q-Tof Mass Spectrometries and Molecular Cloning. General and Comparative Endocrinology, 118, 161–172.

144. Yasuda-Kamatani, Y. and Yasuda, A. (2006) Characteristic expression patterns of allatostatin-like peptide, FMRFamide-related peptide, orcokinin, tachykinin-related peptide, and SIFamide in the olfactory system of crayfishProcambarus clarkii. The Journal of Comparative Neurology, 496, 135–147.

145. Yasuda-Kamatani, Y. and Yasuda, A. (2004) APSGFLGMRamide is a unique tachykinin-related peptide in crustaceans. European Journal of Biochemistry, 271, 1546–1556.

146. Ye, H., Hui, L., Kellersberger, K. and Li, L. (2013) Mapping of neuropeptides in the crustacean stomatogastric nervous system by imaging mass spectrometry. Journal of the American Society for Mass Spectrometry, 24, 134–147.

147. Ye, H., Wang, J., Zhang, Z., Jia, C., Schmerberg, C., Catherman, A.D., Thomas, P.M., Kelleher, N.L. and Li, L. (2015) Defining the Neuropeptidome of the Spiny Lobster Panulirus interruptus Brain Using a Multidimensional Mass Spectrometry-Based Platform. Journal of Proteome Research, 14, 4776–4791.

148. Yin, G.-L., Yang, J.-S., Cao, J.-X. and Yang, W.-J. (2006) Molecular cloning and characterization of FGLamide allatostatin gene from the prawn, Macrobrachium rosenbergii. Peptides, 27, 1241–1250.

149. Zhang, Y., Buchberger, A., Muthuvel, G. and Li, L. (2015) Expression and distribution of neuropeptides in the nervous system of the crab Carcinus maenas and their roles in environmental stress. PROTEOMICS, 15, 3969–3979.

150. Zhang, Y., DeLaney, K., Hui, L., Wang, J., Sturm, R.M. and Li, L. (2018) A Multifaceted Mass Spectrometric Method to Probe Feeding Related Neuropeptide Changes in Callinectes sapidus and Carcinus maenas. Journal of the American Society for Mass Spectrometry, 29, 948–960.

151. Zhang, Z., Jia, C. and Li, L. (2012) Neuropeptide analysis with liquid chromatography-capillary electrophoresis-mass spectrometric imaging. Journal of Separation Science, 35, 1779–1784.

152. Zhang, Z., Wang, J., Hui, L. and Li, L. (2011) Membrane-assisted capillary isoelectric focusing coupling with matrix-assisted laser desorption/ionization-Fourier transform mass spectrometry for neuropeptide analysis. Journal of Chromatography A, 1218, 5336–5343.

153. Zhang, Z., Ye, H., Wang, J., Hui, L. and Li, L. (2012) Pressure-Assisted Capillary Electrophoresis Coupling with Matrix-Assisted Laser Desorption/Ionization-Mass Spectrometric Imaging for Quantitative Analysis of Complex Peptide Mixtures. Analytical Chemistry, 84, 7684–7691.

154. Sithigorngul, W., Jaideechoey, S., Saraithongkum, W., Longyant, S. and Sithigorngul, P. (1999) Purification and characterization of an isoform of crustacean hyperglycemic hormone from the eyestalk of. J Exp Zool, 284, 217–224.

155. Chen, R., Jiang, X., Prieto Conaway, M.C., Mohtashemi, I., Hui, L., Viner, R. and Li, L. (2010) Mass Spectral Analysis of Neuropeptide Expression and Distribution in the Nervous System of the Lobster Homarus americanus. Journal of Proteome Research, 9, 818–832.

156. Cock, P.J.A., Antao, T., Chang, J.T., Chapman, B.A., Cox, C.J., Dalke, A., Friedberg, I., Hamelryck, T., Kauff, F., Wilczynski, B. et al. (2009) Biopython: freely available Python tools for computational molecular biology and bioinformatics. Bioinformatics, 25, 1422–1423.

157. Ikai, A. (1980) Thermostability and Aliphatic Index of Globular-Proteins. J Biochem-Tokyo, 88, 1895–1898.

158. Boman, H.G. (2003) Antibacterial peptides: basic facts and emerging concepts. J Intern Med, 254, 197–215.

159. Abramson, J., Adler, J., Dunger, J., Evans, R., Green, T., Pritzel, A., Ronneberger, O., Willmore, L., Ballard, A.J., Bambrick, J. et al. (2024) Accurate structure prediction of biomolecular interactions with AlphaFold 3. Nature, 630, 493–500.

160. Lin, Z., Akin, H., Rao, R., Hie, B., Zhu, Z., Lu, W., Smetanin, N., Verkuil, R., Kabeli, O., Shmueli, Y. et al. (2023) Evolutionary-scale prediction of atomic-level protein structure with a language model. Science, 379, 1123–1130.

161. Fields, L., Vu, N.Q., Dang, T.C., Yen, H.C., Ma, M., Wu, W., Gray, M. and Li, L. (2024) EndoGenius: Optimized Neuropeptide Identification from Mass Spectrometry Datasets. J Proteome Res, 23, 3041–3051.

162. Dang, T.C., Fields, L. and Li, L. (2024) MotifQuest: An Automated Pipeline for Motif Database Creation to Improve Peptidomics Database Searching Programs. J Am Soc Mass Spectrom, 35, 1902–1912.

163. Fields, L., Ma, M., Delaney, K., Phetsanthad, A. and Li, L.J. (2024) A crustacean neuropeptide spectral library for data-independent acquisition (DIA) mass spectrometry applications. Proteomics, 24.

164. Vu, N.Q., Yen, H.C., Fields, L., Cao, W.F. and Li, L.J. (2023) HyPep: An Open-Source Software for Identification and Discovery of Neuropeptides Using Sequence Homology Search. Journal of Proteome Research.

165. DeLaney, K., Cao, W.F., Ma, Y.D., Ma, M.M., Zhang, Y.Z. and Li, L.J. (2020) PRESnovo: Prescreening Prior to Sequencing to Improve Accuracy and Sensitivity of Neuropeptide Identification. Journal of the American Society for Mass Spectrometry, 31, 1358–1371.

166. Lin, Z.M., Akin, H., Rao, R.S., Hie, B., Zhu, Z.K., Lu, W.T., Smetanin, N., Verkuil, R., Kabeli, O., Shmueli, Y. et al. (2023) Evolutionary-scale prediction of atomic-level protein structure with a language model. Science, 379, 1123–1130.

167. Wang, L., Zeng, Z., Xue, Z. and Wang, Y. (2024) DeepNeuropePred: A robust and universal tool to predict cleavage sites from neuropeptide precursors by protein language model. Comput Struct Biotechnol J, 23, 309–315.

168. Wilkinson, R.E., Kraichely, K.N., Hendy, C.M., Buchanan, L.E., Parnham, S. and Giuliano, M.W. (2022) The neuropeptide galanin adopts an irregular secondary structure. Biochem Biophys Res Commun, 626, 121–128.

169. Dickinson, P.S., Samuel, H.M., Stemmler, E.A. and Christie, A.E. (2019) SIFamide peptides modulate cardiac activity differently in two species of Cancer crab. Gen Comp Endocrinol, 282, 113204.

170. Dickinson, P.S. and Powell, D.J. (2023) Diversity of neuropeptidergic modulation in decapod crustacean cardiac and feeding systems. Curr Opin Neurobiol, 83, 102802.

171. Muscato, A.J., Walsh, P., Pong, S., Pupo, A., Gross, R.J., Christie, A.E., Hull, J.J. and Dickinson, P.S. (2021) Does Differential Receptor Distribution Underlie Variable Responses to a Neuropeptide in the Lobster Cardiac System? Int J Mol Sci, 22, 8703–8703.

172. Xu, Z., Wei, Y., Guo, S., Lin, D. and Ye, H. (2020) Short neuropeptide F enhances the immune response in the hepatopancreas of mud crab (Scylla paramamosain). Fish Shellfish Immunol, 101, 244–251.

173. Zuo, H., Yuan, J., Niu, S., Yang, L., Weng, S., He, J. and Xu, X. (2018) A molting-inhibiting hormone-like protein from Pacific white shrimp Litopenaeus vannamei is involved in immune responses. Fish Shellfish Immunol, 72, 544–551.

174. Christie, A.E., Yu, A., Roncalli, V., Pascual, M.G., Cieslak, M.C., Warner, A.N., Lameyer, T.J., Stanhope, M.E., Dickinson, P.S. and Joe Hull, J. (2018) Molecular evidence for an intrinsic circadian pacemaker in the cardiac ganglion of the American lobster, <i>Homarus americanus</i> - Is diel cycling of heartbeat frequency controlled by a peripheral clock system? Marine Genomics, 41, 19–30.

175. Harzsch, S., Dircksen, H. and Beltz, B.S. (2009) Development of pigment-dispersing hormone-immunoreactive neurons in the American lobster: homology to the insect circadian pacemaker system? Cell Tissue Res, 335, 417–429.

176. Strauss, J. and Dircksen, H. (2010) Circadian clocks in crustaceans: identified neuronal and cellular systems. Front Biosci (Landmark Ed), 15, 1040–1074.

177. Lehtonen, M.P. and Burnett, L.E. (2016) Effects of Hypoxia and Hypercapnic Hypoxia on Oxygen Transport and Acid-Base Status in the Atlantic Blue Crab, Callinectes sapidus, During Exercise. J Exp Zool A Ecol Genet Physiol, 325, 598–609.

178. Delorenzi, A., Dimant, B., Frenkel, L., Nahmod, V.E., Nassel, D.R. and Maldonado, H. (2000) High environmental salinity induces memory enhancement and increases levels of brain angiotensin-like peptides in the crab Chasmagnathus granulatus. J Exp Biol, 203, 3369–3379.

179. Sun, S., Zhu, M., Pan, F., Feng, J. and Li, J. (2020) Identifying Neuropeptide and G Protein-Coupled Receptors of Juvenile Oriental River Prawn (Macrobrachium nipponense) in Response to Salinity Acclimation. Front Endocrinol (Lausanne), 11, 623.

180. Gong, J., Yu, K., Shu, L., Ye, H.H., Li, S.J. and Zeng, C.S. (2015) Evaluating the effects of temperature, salinity, starvation and autotomy on molting success, molting interval and expression of ecdysone receptor in early juvenile mud crabs,. J Exp Mar Biol Ecol, 464, 11–17.

181. Sauer, C.S. and Li, L. (2021) Mass Spectrometric Profiling of Neuropeptides in Response to Copper Toxicity via Isobaric Tagging. Chemical Research in Toxicology, 34, 1329–1336.

182. Pulver, S.R. and Marder, E. (2002) Neuromodulatory complement of the pericardial organs in the embryonic lobster, Homarus americanus. J Comp Neurol, 451, 79–90.

183. Jiang, X., Chen, R., Wang, J., Metzler, A., Tlusty, M. and Li, L. (2012) Mass Spectral Charting of Neuropeptidomic Expression in the Stomatogastric Ganglion at Multiple Developmental Stages of the Lobster Homarus americanus. ACS Chemical Neuroscience, 3, 439–450.

184. Hu, M., Helfenbein, K., Buchberger, A.R., DeLaney, K., Liu, Y. and Li, L. (2021) Exploring the Sexual Dimorphism of Crustacean Neuropeptide Expression Using Callinectes sapidus as a Model Organism. J Proteome Res, 20, 2739–2750.

185. Ao, C.M., Shi, L.L., Wang, W., Wang, C.G. and Chan, S.F. (2022) Neuroparsin 1 (MrNP1) and Neuroparsin 2 (MrNP2) Are Involved in the Regulation of Vitellogenesis in the Shrimp. Front Mar Sci, 9.

186. Cui, Z., Liu, Y., Yuan, J., Zhang, X., Ventura, T., Ma, K.Y., Sun, S., Song, C., Zhan, D., Yang, Y. et al. (2021) The Chinese mitten crab genome provides insights into adaptive plasticity and developmental regulation. Nat Commun, 12, 2395.

187. Gutekunst, J., Andriantsoa, R., Falckenhayn, C., Hanna, K., Stein, W., Rasamy, J. and Lyko, F. (2018) Clonal genome evolution and rapid invasive spread of the marbled crayfish. Nat Ecol Evol, 2, 567–573.

188. Stein, W., DeMaegd, M.L., Benson, A.M., Roy, R.S. and Vidal-Gadea, A.G. (2022) Combining Old and New Tricks: The Study of Genes, Neurons, and Behavior in Crayfish. Front Physiol, 13, 947598.

189. Veldsman, W.P., Ma, K.Y., Hui, J.H.L., Chan, T.F., Baeza, J.A., Qin, J. and Chu, K.H. (2021) Comparative genomics of the coconut crab and other decapod crustaceans: exploring the molecular basis of terrestrial adaptation. BMC Genomics, 22, 313.

190. Schneider, A.C., Seichter, H.A., Neupert, S., Hochhaus, A.M. and Smarandache-Wellmann, C.R. (2018) Profiling neurotransmitters in a crustacean neural circuit for locomotion. PLoS One, 13, e0197781.

191. Cao, Q., Wang, Y., Chen, B., Ma, F., Hao, L., Li, G., Ouyang, C. and Li, L. (2019) Visualization and Identification of Neurotransmitters in Crustacean Brain via Multifaceted Mass Spectrometric Approaches. ACS Chem Neurosci, 10, 1222–1229.

192. Kravitz, E.A. and Sengupta, S. (2024) Crustaceans played a primary role in establishing gamma-aminobutyric acid as a neurotransmitter. Curr Opin Insect Sci, 65, 101252.

193. Hedrich, U.B. and Stein, W. (2008) Characterization of a descending pathway: activation and effects on motor patterns in the brachyuran crustacean stomatogastric nervous system. J Exp Biol, 211, 2624–2637.

194. Rue, M.C.P., Alonso, L.M. and Marder, E. (2022) Repeated applications of high potassium elicit long-term changes in a motor circuit from the crab, Cancer borealis. iScience, 25, 104919.

195. Stein, W. and Stadele, C. (2024) Neuromodulator-induced temperature robustness in a motor pattern: a comparative study between two decapod crustaceans. J Exp Biol, 227.

196. Tian, Z. and Jiao, C. (2019) Molt-dependent transcriptome analysis of claw muscles in Chinese mitten crab Eriocheir sinensis. Genes Genomics, 41, 515–528.

197. Bostjancic, L.L., Francesconi, C., Rutz, C., Hoffbeck, L., Poidevin, L., Kress, A., Jussila, J., Makkonen, J., Feldmeyer, B., Balint, M. et al. (2022) Host-pathogen coevolution drives innate immune response to Aphanomyces astaci infection in freshwater crayfish: transcriptomic evidence. BMC Genomics, 23, 600.

198. Vu, N.Q., DeLaney, K. and Li, L. (2021) Neuropeptidomics: Improvements in Mass Spectrometry Imaging Analysis and Recent Advancements. Current protein & peptide science, 22, 158–169.

